# Whole genome screen reveals a novel relationship between *Wolbachia* levels and *Drosophila* host translation

**DOI:** 10.1101/380485

**Authors:** Yolande Grobler, Chi Y. Yun, David J. Kahler, Casey M. Bergman, Hangnoh Lee, Brian Oliver, Ruth Lehmann

## Abstract

*Wolbachia* is an intracellular bacterium that infects a remarkable range of insect hosts. Insects such as mosquitos act as vectors for many devastating human viruses such as Dengue, West Nile, and Zika. Remarkably, *Wolbachia* infection provides insect hosts with resistance to many arboviruses thereby rendering the insects ineffective as vectors. To utilize *Wolbachia* effectively as a tool against vector-borne viruses a better understanding of the host-*Wolbachia* relationship is needed. To investigate *Wolbachia*-insect interactions we used the *Wolbachia/Drosophila* model that provides a genetically tractable system for studying host-pathogen interactions. We coupled genome-wide RNAi screening with a novel high-throughput fluorescence in situ hybridization (FISH) assay to detect changes in *Wolbachia* levels in a *Wolbachia*-infected *Drosophila* cell line JW18. 1117 genes altered *Wolbachia* levels when knocked down by RNAi of which 329 genes increased and 788 genes decreased the level of *Wolbachia.* Validation of hits included in depth secondary screening using *in vitro* RNAi, *Drosophila* mutants, and *Wolbachia*-detection by DNA qPCR. A diverse set of host gene networks was identified to regulate *Wolbachia* levels and unexpectedly revealed that perturbations of host translation components such as the ribosome and translation initiation factors results in increased *Wolbachia* levels both *in vitro* using RNAi and *in vivo* using mutants and a chemical-based translation inhibition assay. This work provides evidence for *Wolbachia*-host translation interaction and strengthens our general understanding of the *Wolbachia*-host intracellular relationship.

**Author summary:** Insects such as mosquitos act as vectors to spread devastating human diseases such as Dengue, West Nile, and Zika. It is critical to develop control strategies to prevent the transmission of these diseases to human populations. A novel strategy takes advantage of an endosymbiotic bacterium *Wolbachia pipientis.* The presence of this bacterium in insect vectors prevents successful transmission of RNA viruses. The degree to which viruses are blocked by *Wolbachia* is dependent on the levels of the bacteria present in the host such that higher *Wolbachia* levels induce a stronger antiviral effect. In order to use *Wolbachia* as a tool against vector-borne virus transmission a better understanding of host influences on *Wolbachia* levels is needed. Here we performed a genome-wide RNAi screen in a model host system *Drosophila melanogaster* infected with *Wolbachia* to identify host systems that affect *Wolbachia* levels. We found that host translation can influence *Wolbachia* levels in the host.

## Introduction

Insects are common vectors for devastating human viruses such as Zika, Yellow Fever, and Dengue. A novel preventative strategy has emerged to combat vector-borne diseases that exploits the consequences of vector-insect infection with the bacteria *Wolbachia pipientis* [1–4]. *Wolbachia* is a vertically transmitted, gram-negative intracellular bacterium known to infect 40-70% of all insects [5, 6]. *Wolbachia* provides hosts with resistance to pathogens such as viruses [7–10]. Remarkably, *Wolbachia* infections can reduce host viral load enough to render insect hosts incapable of transmitting disease-causing viruses effectively [1, 2, 11–24]. The relationship between *Wolbachia* and a host is complex and dynamic. Understanding how bacterial levels can change is vital because it dictates how *Wolbachia* manipulates the host insect. For example, the antiviral protection provided by *Wolbachia* is strongest when *Wolbachia* levels within a host are high [10, 25–27]. On the other hand, *Wolbachia* can become deleterious to the host when *Wolbachia* population levels are too high leading to cellular damage and reduced lifespan[28–30]. To apply *Wolbachia* as an effective tool to combat vector-borne viruses we need a better understanding of host influences on *Wolbachia* levels.

*Wolbachia* infects a large host and tissue range suggesting interaction with various host systems and pathways for successful intracellular maintenance within a host [5, 31]. To date, reports suggest that *Wolbachia* levels may be influenced in various contexts by interaction with host cytoskeletal components [32–35], the host ubiquitin/proteasome [36], host autophagy [37], and by host miRNAs [16, 38]. A comprehensive analysis of host systems that influence *Wolbachia* levels has not been carried out and will further our knowledge of this symbiotic relationship and reveal molecular mechanisms that occur between *Wolbachia* and the host to maintain it.

*Wolbachia-host* interactions can be studied in the genetically tractable *Drosophila melanogaster* system which allows for the systematic dissection of host signaling pathways that interact with the bacteria using the wide array of genetic and genomic tools available. The *Drosophila* system enables rapid unbiased screening of host factors that impact *Wolbachia* at the cellular and organismal level. While some influences on the relationship, such as systemic effects, require studies in the whole organism, many aspects of molecular and cellular signaling can be studied in a *Drosophila* cell culture-based system. *Drosophila* cells are particularly amenable to genome-scale screens because of the ease and efficacy of RNAi in this system [39]-[40]. Previous cell culture-based RNAi screening has been a successful approach to study a wide range of intracellular bacteria-host interactions in *Drosophila* cell lines [41–44]. Thus, we reasoned that this was a feasible approach for studying *Wolbachia-host* interactions.

Here we performed a whole genome RNAi screen in a *Wolbachia-infected Drosophila* cell line, JW18, which was originally derived from *Wolbachia-infected Drosophila* embryos and has previously proven suitable for high-throughput assays [36, 45]. The goal was to determine in an unbiased and comprehensive manner which host systems affect intracellular *Wolbachia* levels. The primary screen identified 1117 host genes that robustly altered *Wolbachia* levels. Knock down of 329 of these genes resulted in increased *Wolbachia* levels whereas 788 genes led to decreased *Wolbachia* levels. To characterize these genes, we generated manually curated categories, performed Gene Ontology enrichment analysis, and identified enriched host networks using bioinformatic analysis tools. The effects on *Wolbachia* levels were validated in follow-up RNAi assays that confirmed *Wolbachia* changes visually by RNA FISH as well as quantitatively using a highly sensitive DNA qPCR assay. We uncovered an unexpected role of host translation components such as the ribosome and translation initiation factors in suppressing *Wolbachia* levels both in tissue culture using RNAi and in the fly using mutants and a chemical-based translation inhibition assay. Furthermore, we show an inverse relationship between *Wolbachia* levels and host translation levels in JW18 cells as well as in a characterized *Wolbachia* niche, namely the *Drosophila* testis hub. This work provides strong evidence for a relationship between *Wolbachia* and host translation and strengthens our general understanding of the *Wolbachia-host* intracellular relationship.

## Results

### Characterization of *Wolbachia-infected* JW18 *Drosophila* cells

*Wolbachia* is an intracellular bacterium that resides within a wide range of insect hosts. To identify host factors that enhance or suppress intracellular *Wolbachia* levels, we performed a genome-wide RNAi screen in *Wolbachia-infected* JW18 *Drosophila* cells that were originally derived from *Wolbachia-infected* embryos [45]. In order to visually detect *Wolbachia* levels we established a specific and sensitive RNA Fluorescence In Situ Hybridization (FISH) method consisting of a set of 48 fluorescently labeled DNA oligos that collectively bind in series to the target *Wolbachia 23s rRNA* (Fig 1A). This enabled detection of infection levels ranging from as low as a single bacterium in a cell to a highly infected cell and could clearly distinguish *Wolbachia-infected* cells from *Wolbachia-free* cells (Fig 1B). Thus, we were able to assess *Wolbachia* infection levels in the JW18 cell population and found that under our culturing conditions we could stably maintain JW18 cells with a Wolbachia infection level of 14% of the JW18 cells (Fig 1C). Of the infected cells, 73% of the cells had a low *Wolbachia* infection (1-10 bacteria), 13.5% had a medium infection (11-30 bacteria), and 13.5% were highly infected (>30 bacteria). These experiments confirmed the feasibility and sensitivity of RNA FISH to detect different levels of *Wolbachia* infection in *Drosophila* cells in a highly sensitive manner.

**Figure 1.**
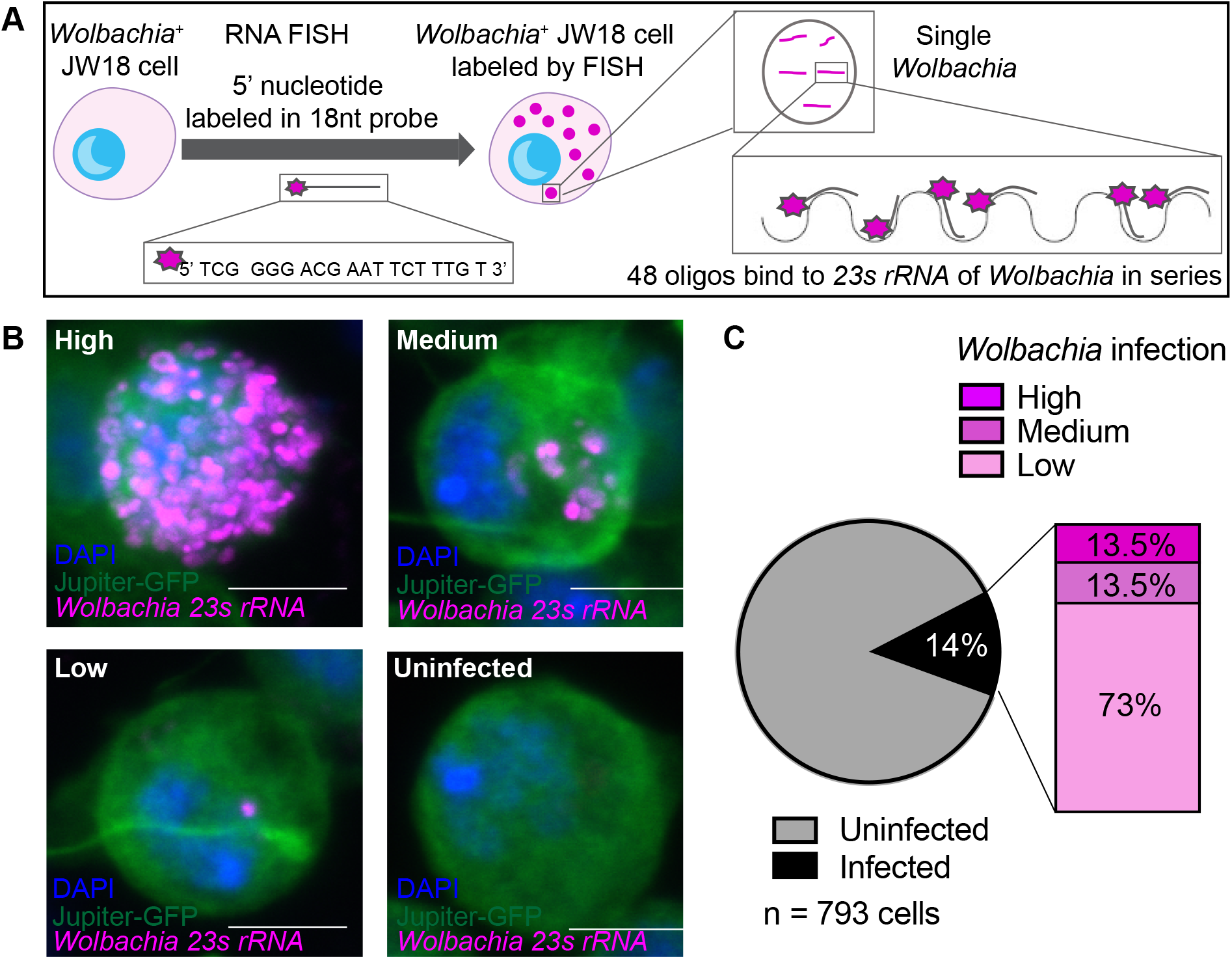
Visualization of infection dynamics in the JW18 cell line. **(A)** Schematic of *Wolbachia* detection by RNA Fluorescent In Situ Hybridization (FISH) using a sensitive and specific set of 48 5’-fluorescently-labeled oligos that bind in series to the *23s rRNA* of the *Wolbachia* within a host cell. **(B)** *Wolbachia-infected* JW18 cells labeled by *23s rRNA* FISH probe can detect different infection levels in a highly specific manner. Scale bar 5μm. **(C)** *Wolbachia* infection within the JW18 population is steadily maintained at 14% of the total cells in the population. Of the *Wolbachia* infected cells, the majority (73%) of cells have a low *Wolbachia* infection (1-10 bacteria per cell), 13.5% contain a medium *Wolbachia* infection (11-30 bacteria), and 13.5% of the infected JW18 cells have a high infection level (>30 bacteria) (n=793 cells).

Prior to screening we characterized the JW18 cell line and its associated *Wolbachia* strain by generating a JW18 DNA library and sequencing it using DNAseq technology (S1 Fig). This allowed for phylogenetic analysis of the *Wolbachia* strain and revealed that it clustered most closely with the avirulent wMel strain which is well characterized for its antiviral effect on RNA-based viruses in *Drosophila* as well as in mosquitos (S1A Fig) [1, 2, 7, 8]. Further analysis included gene copy number variation of the *Wolbachia* genome and identified one deleted and one highly duplicated region (3-4 fold increased) (S1B Fig). The deleted region contained eight genes known as the “Octomom” region postulated to influence virulence [27, 46]. The loss of “Octomom” has also been reported in wMelPop-infected mosquito cell lines after extended passaging over 44 months [47]. This suggests that loss of this region happened independently in two cases and may be related to passage in cell culture. A highly duplicated region spans approximately from positions 91,800-127,100 and contains 38 full or partial genes, including those with unknown function as well as genes predicted to be involved in metabolite synthesis and transport, molecular chaperones, DNA polymerase III subunit, DNA gyrase subunit, and 50S ribosomal subunits. For analysis of gene copy number variation in the JW18 cell line, the DNA library was aligned to the Release 6 reference genome of *D. melanogaster.* This revealed that the cell line is of male origin with an X:A chromosomal ratio of 1:2 and tetraploid in copy number (S1C Fig). Bioinformatic analysis on genes of high or low copy number did not reveal an enrichment for any particular molecular or cellular functional class and the majority (72%) of genes in the JW18 cell line were at copy numbers expected for a tetraploid male genotype (4 copies on autosomes, 2 copies on X). This made the JW18 cell line suitable for RNAi screening.

As a first step to uncovering *Wolbachia-host* interactions, we asked whether gene expression changes occur in the host during stable *Wolbachia* infection. To do this, a control *Wolbachia-free* version of the JW18 cell line was generated through doxycycline treatment (JW18TET) (S2A Fig). A comparison of host gene expression changes in the presence and absence of *Wolbachia* through RNAseq analysis revealed 308 and 559 host genes that were up-or down-regulated respectively by two-fold or more (padj<0.05) (S2B Fig). Of these genes, 21 displayed major expression changes of log2 fold >4 (S2C Fig). The presence of *Wolbachia* led to elevated gene expression of several components of the host immune response including the Gram-negative antimicrobial peptide *Diptericin B (DptB),* which was the most highly upregulated gene in the presence of *Wolbachia* (S2C Fig). Gene ontology (GO) analysis further confirmed a host immune response with enriched terms such as ‘response to other organism’ and ‘peptidoglycan binding’ that included genes for antimicrobial peptides (attA, AttB, AttC, DptB, LysB) and peptidoglycan receptors (PGRP-SA, −SD, −LB, −LF) as well as antioxidants such as Jafrac2, Prx2540-1, Prx2540-2, Pxn, GstS1 with ‘peroxiredoxin’ and ‘peroxidase activity’. Other expression changes included extracellular matrix components such as upregulation of collagen type IV *(Col4a1* and *vkg)* and downregulation of genes for integral components of the plasma membrane including cell adhesion components *(kek5, mew, Integrin,* and *tetraspanin 42Ed* and *39D).* Gene ontology analysis further identified a significant enrichment of ion transporters and channels that were downregulated as well as genes encoding several proteins such as myosin II, projectin and others associated with the muscle Z-disc that were downregulated. Finally, we observed an overall upregulation of host proteasome components at the RNA level in the presence of *Wolbachia* (S9 Fig), which is in line with proteomics data of proteasome upregulation in the presence of *Wolbachia* [48, 49]. In summary, these host factors may play an important role in the *Wolbachia-host* relationship however their specific roles in this interaction remain to be determined.

### Genome-wide RNAi screen to identify host genes that affect *Wolbachia* levels within the host cell

The screening approach combined the visual RNA FISH *Wolbachia* detection assay (Fig 1) with *in vitro* RNAi knockdown of host genes to ask which host genes influence *Wolbachia* levels (Fig 2A). Prior to screening, we tested whether RNAi was a feasible approach in JW18 cells. First, we confirmed that RNAi had no adverse effects such as cytotoxicity on the cells using a negative control dsRNA targeting LacZ which was not present in our system (Fig 2B,C). Second, we tested RNAi knockdown efficiency in the JW18 cell line. To do this a Jupiter-GFP transgene present in the cell line was targeted for knockdown using dsRNA to GFP. High knockdown efficiency was achieved using this RNAi protocol as seen by the efficient knockdown of the Jupiter-GFP transgene both visually by RNA FISH (Fig 2D) and by protein levels as shown by Western blot (Fig 2E) compared with either the no knockdown (Fig 2B) or LacZ knockdown (Fig 2C) conditions. This confirmed the suitability of the JW18 cell line for an RNAi-based screening approach. For controls that alter *Wolbachia* levels, we identified a host ribosomal gene, *RpL40,* from a pilot screen that consistently led to increased *Wolbachia* levels when depleted by RNAi (Fig 2F) compared to cells that were not treated by RNAi (Fig 2B) or treated with *lacZ* dsRNA treatment (Fig 2C). We achieved 96.3% RNAi knockdown efficiency as confirmed by qPCR for *RpL40* levels relative to a no knockdown control (Fig 2G). At the time of the screen we did not know of any host protein whose knockdown would decrease *Wolbachia* levels. Therefore, as a Wolbachia-decreasing control, cells were incubated with 5μM doxycycline for 5 days which successfully reduced the *Wolbachia* levels in the cells below a detectable level by RNA FISH (Fig 2H). To quantify the effect of the controls on *Wolbachia* levels we isolated genomic DNA from each treated sample and used quantitative PCR DNA amplification to detect the number of *Wolbachia* genomes per cell by measuring *Wolbachia* wspB copy number relative to the *Drosophila* gene RpL11 (Fig 2I). Relative to control cells, the RNAi treatment with RpL40 resulted in a 3.4-fold increase in *Wolbachia,* doxycycline decreased *Wolbachia* levels, whereas LacZ and GFP RNAi had no significant effect confirming that our controls allowed us to manipulate *Wolbachia* levels in the JW18 cell line and that this cell line with its relative low infection rate (Fig 1C) provided a sensitive tool for detecting dynamic changes in *Wolbachia* levels through an RNAi screening approach.

**Figure 2.**
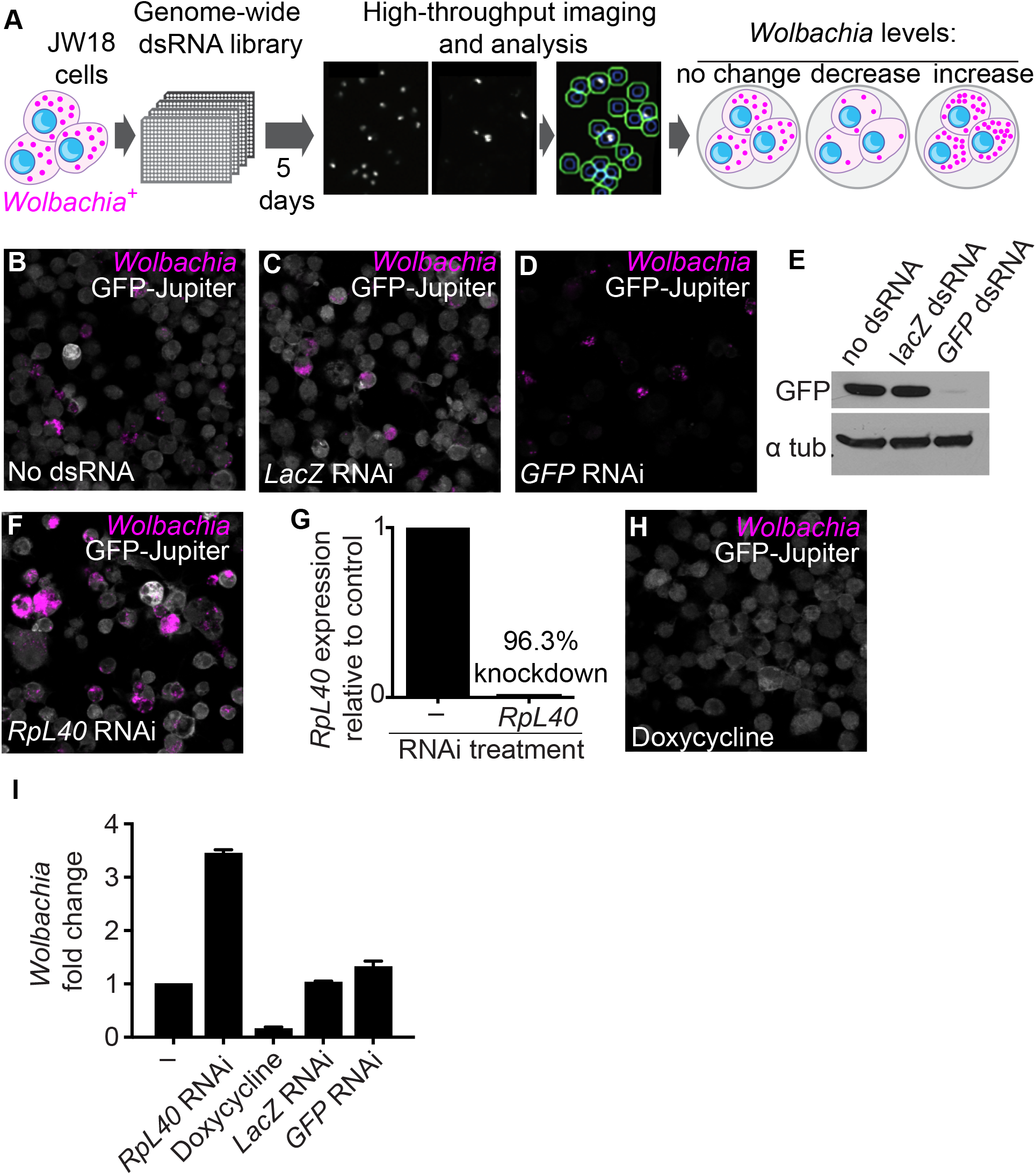
Genome-wide screening approach to find novel *Wolbachia-host* interactions in *Wolbachia-infected* JW18 *Drosophila* cells. **(A)** Schematic of screen layout. *Wolbachia-infected* JW18 cells are seeded into 384-well plates pre-arrayed with the DRSC version 2.0 whole genome RNAi library designed to include dsRNAs that target the whole *Drosophila* genome. All plates were screened in triplicate. Cells and dsRNA were incubated for 5 days before processing for an automated high-throughput RNA-FISH assay to detect changes in *Wolbachia 23s rRNA* levels. **(B)** Representative image of *Wolbachia* detection at 20x with the *23s rRNA Wolbachia* FISH probe (magenta) in JW18 cells containing a *GFP-Jupiter* transgene labeling microtubules (grey). **(C)** Negative control dsRNA against *LacZ* not present in our system. **(D, E)** RNAi control against *GFP-Jupiter* shows efficient knockdown visually as well as by protein levels. **(F, G)** Positive control for increasing *Wolbachia* levels using efficient RNAi-mediated silencing of host gene *RpL40.* **(H)** Positive control for decreasing *Wolbachia* levels through treatment with doxycycline for 5 days. **(I)** Quantification of *Wolbachia* level fold-change relative to untreated JW18 cells using DNA qPCR.

The layout of the whole genome screen is illustrated in Fig 2A. Briefly, Wolbachia-infected JW18 cells were incubated with the DRSC *Drosophila* Whole Genome RNAi Library version 2.0 which was pre-arrayed in 384 well tissue culture plates such that each well contained a specific dsRNA amplicon to target one host gene. The 5-day incubation period allowed for efficient host gene knockdown. Thereafter the cells were processed for RNA FISH detection of *Wolbachia 23s rRNA.* Total fluorescence signal was detected using automated microscopy and served as a readout for *Wolbachia* levels within each plate well. Host cells within each well were detected by DAPI staining. Finally, the *Wolbachia* fluorescence signal was divided by the total number of DAPI-stained host cells detected to provide an average *Wolbachia* per cell readout which was normalized to the plate average (represented as a robust Z score). The library was screened in triplicate.

The raw screening data was subjected to several quality control steps (S3 Fig). Briefly, we realigned the DRSC Version 2.0 Whole Genome RNAi library dsRNA amplicons with Release 6 of the *D. melanogaster* genome using the bioinformatic tool UP-TORR [50]. This provided an accurate updated description of the gene target for each dsRNA amplicon. Initially the library included 24 036 unique dsRNA amplicons targeting 15 589 genes, however owing to updates in gene organization and annotation models of the reference genome since the initial release of the library we removed 1499 outdated amplicons from our subsequent analysis as they were no longer predicted to have gene targets (S1 Table). We also excluded 1481 amplicons that were annotated in UP-TORR to target multiple genes (S2 and S3 Tables). We further excluded 66 amplicons for a positional effect on the dsRNA library tissue culture plates at the A1 position (S4 Fig, S4 Table). Thus, we effectively screened 20 990 unique dsRNA amplicons targeting 14 024 genes (80% of *D. melanogaster* Release 6 genome). A further quality control step to reduce false positive hits was to cross-reference potential hits with RNAseq gene expression data for the JW18 cell line to exclude genes with undetectable expression in the cell line (S5 Table).

To identify and select for hits from the primary data, we first analyzed the screen-wide controls. A plot of all controls included in the whole genome screen revealed that *RpL40* knockdown increased *Wolbachia* levels (median robust Z score of 2.2), conversely doxycycline treatment decreased *Wolbachia* levels throughout the screen (median robust Z of −3.5), whereas a standard control included in the whole genome library, *Rho1,* and *GFP* RNAi knockdown did not significantly affect *Wolbachia* levels (Fig 3A). We used this range as a guide to set robust Z limits for primary hits at ≥ 1.5 or ≤ −1.5. Every dsRNA amplicon was screened in triplicate. To be considered as a ‘hit’ amplicon at least 2 of the 3 replicates needed to have significant robust Z scores (S3 Fig). To categorize the primary screen hits, each gene was assigned to a ‘High’, ‘Medium’, and “Low’ bin based on the confidence level (S3 Fig). This was determined based on the total number of different dsRNA amplicons representing a hit gene in the library and how many of these dsRNA amplicons had a significant effect on *Wolbachia* levels (S3 Fig). In this manner, we were able to stratify the primary screen hits to assist in follow up analysis.

**Figure 3.**
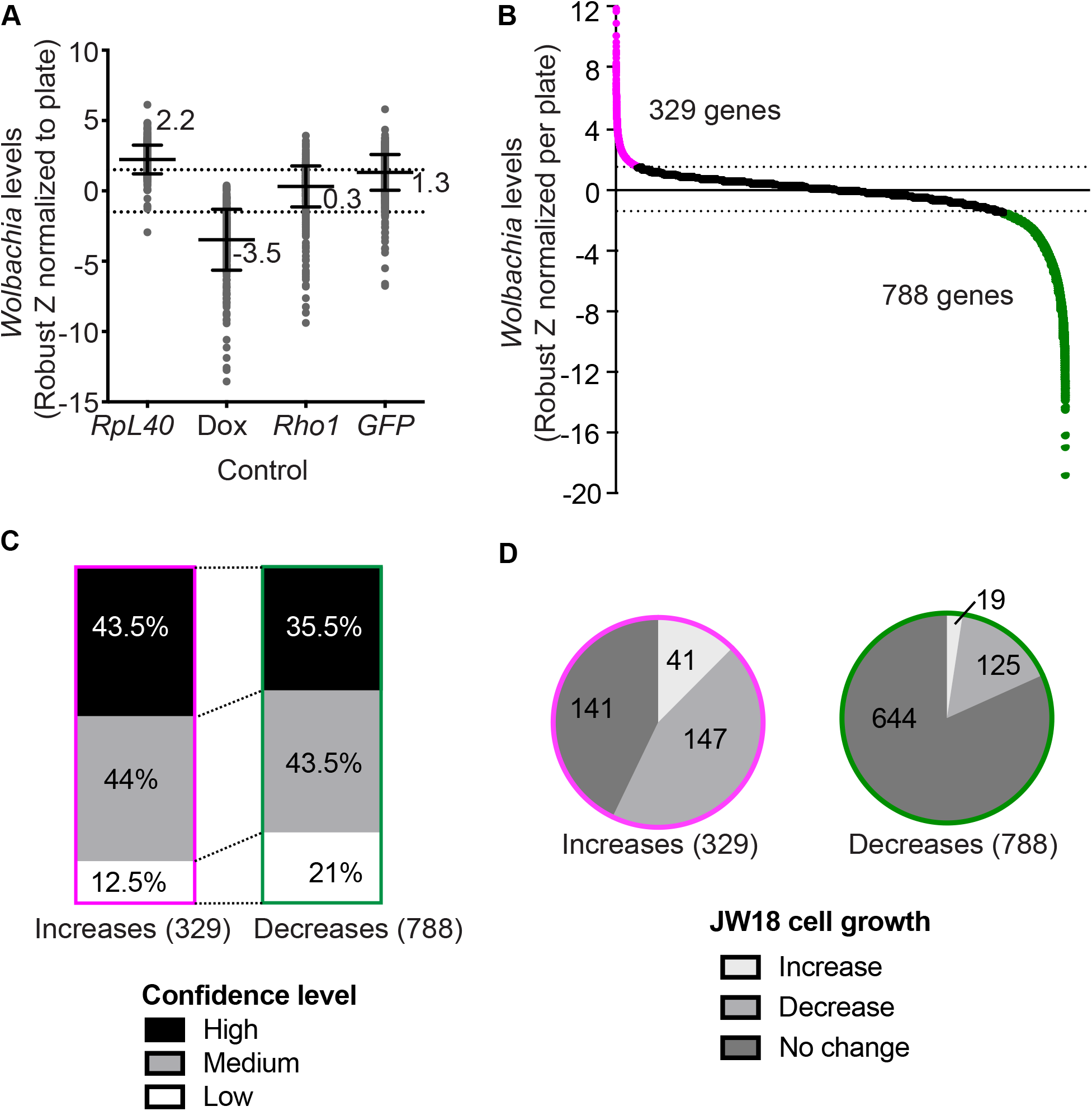
Genome-wide screen controls and primary results. **(A)** Plot of screen-wide controls’ effect on *Wolbachia* levels (Robust Z score normalized for each plate) included RNAi silencing of a host ribosomal gene *RpL40* to increase *Wolbachia,* doxycycline treatment to decrease *Wolbachia,* RNAi silencing of *Rho1* as a negative control, and RNAi silencing of a *GFP-Jupiter* transgene as a positive control for RNAi in our system that has no effect on *Wolbachia* levels. Bars represent the median and interquartile range of the robust Z scores for each control in the whole genome screen. **(B)** The whole genome screen yielded 1117 primary hits with a robust Z score of ≥1.5 or ≤-1.5. These genes included 329 genes that increase *Wolbachia* (magenta) and 788 genes that decrease *Wolbachia* (green) upon RNAi knockdown. **(C)** 1117 primary hits categorized according to confidence level (see **S3 Fig**). **(D)** 1117 primary hits’ effects on JW18 cell growth. For the genes that significantly increased *Wolbachia* levels (magenta), 12% (41 hits) significantly increased cell growth (robust Z > 1), 43% (141 hits) did not have a significant effect on cell growth, and 45% (147 hits) resulted in significant decreases in cell growth (robust Z < −1). For genes that significantly decreased *Wolbachia* levels (green), 2.4% (19 hits) significantly increased cell growth (robust Z > 1), 82% (644 hits) did not have a significant effect on cell growth, and 16% (125 hits) significantly decreased cell growth (robust Z < −1).

### Identification of 1117 host genes that influence *Wolbachia* levels within the host cell

The screen identified 1117 genes that when knocked down had a significant effect on the *Wolbachia* levels in JW18 cells (S6 Table). Knock down of 329 of the 1117 genes resulted in increased *Wolbachia* levels, suggesting that these genes normally restrict *Wolbachia* levels within the host cell (Fig 3B). Knockdown of 788 genes resulted in decreased *Wolbachia* levels, suggesting *Wolbachia* may be dependent on these host genes for survival within the host cell (Fig 3B). For each of the two hit categories, genes were classified by confidence level (described in S3 Fig, and Fig 3C). We found a higher proportion of low confidence hits (21%) in the category of genes that decreased *Wolbachia* levels compared to genes that led to *Wolbachia* level increases which only contained 12.5% low confidence hits. To analyze the expression of the 1117 genes, the hits were distributed into 9 bins based on their gene expression level from JW18 RNAseq data (S5A Fig). Hits displayed a wide range of expression and an enrichment of low expression for hits that decreased *Wolbachia* levels (S5B Fig). We did not observe any biases for variation in gene DNA copy number based on DNAseq data for the JW18 cell line (S5C Fig).

Next, we asked whether changes in *Wolbachia* levels could be explained by effects on host cell growth or were independent of effects on host cell growth. We measured cell growth using the raw screen data by normalizing the number of cells scanned per well (DAPI) to the number of fields of view required to capture the cells. This allowed us to generate a robust Z score measure of cell growth effects for the 1117 genes identified as hits. For genes that increased *Wolbachia* levels, 12% (41 genes) increased cell growth (robust Z>1), 45% (147 genes) decreased cell growth (robust Z<-1), and 43% (141 genes) had no effect on cell growth (Fig 3D, S7 Table). These data suggest that a significant number of gene knockdowns (45%) may indirectly lead to an increase in *Wolbachia* levels through slowed cell growth. However, when we more carefully quantified the degree of *Wolbachia* increase in several of these candidate knockdowns we observed that the Wolbachia level increases were far larger than would be expected from slowed cell growth alone (Fig 5B, discussed in detail later). Importantly, 43% of hits identified had no effect on cell growth whilst increasing Wolbachia levels. For these reasons we argue that the changes in *Wolbachia* levels identified in the screen are not strictly linked to host cell replication. For genes that decreased *Wolbachia* levels, the majority (82%, 644 genes) did not affect cell growth and 2% (19 genes) increased and 16% (125 genes) decreased cell growth (Fig 3D, S7 Table). To summarize, the screen identified 1117 host genes that act to support or suppress *Wolbachia* levels within the host *Drosophila* cell.

### *Wolbachia* suppressors and enhancers function in diverse host pathways and networks

To classify the 1117 gene hits identified in the whole genome screen, we first manually curated the hits using gene annotation available on FlyBase (www.flybase.org) relating to each gene such as gene family, domains, molecular function, gene ontology (GO) information, gene summaries, interactions and pathways, orthologs, and related recent research papers. We identified distinct categories of genes that when knocked down by RNAi increased (Fig 4A) or decreased (Fig 4B) *Wolbachia* levels. The largest gene category that led to decreased *Wolbachia* levels by RNAi knockdown contained genes for host metabolism and transporters suggesting that *Wolbachia* strongly relies on this aspect of the host (Fig 4B). On the other hand, gene knockdowns that increased *Wolbachia* contained many components of the core ribosome network, translation factors, and the core proteasome and regulatory proteins network (Fig 4A). Six of the broad gene categories could be further sub-classified for processes that either enhanced or suppressed *Wolbachia* levels. First, RNAi knockdown of members in the category containing cytoskeleton, cell adhesion and extracellular matrix components leading to decreased *Wolbachia* included cadherins, formins, spectrin and genes involved in microtubule organization, whereas knockdowns that resulted in increased *Wolbachia* were actin and tubulin-related. Second, *Wolbachia* levels may to be sensitive to disturbances in membrane dynamics and trafficking. Specifically, knockdown of SNARE components, endosomal, lysosomal and ESCRT components decreased *Wolbachia,* whereas knockdown of components of COPI, endosome recycling, and several SNAP receptors increased *Wolbachia* levels. Third, disruptions in several cell cycle-related components led to decreased *Wolbachia* levels, however we observed increases in *Wolbachia* levels when we disrupted cytokinesis, the separase complex and the Anaphase Promoting Complex. Fourth, the knockdown of components related to RNA helicases and the exon junction complex decreased *Wolbachia,* while disruption of many spliceosome components led to increased *Wolbachia.* Fifth, epigenetic changes influenced *Wolbachia* levels: knockdown of members involved in heterochromatin silencing, Sin3 complex and coREST led to decreased *Wolbachia* levels, whereas knocking down members of the BRAHMA complex resulted in increased *Wolbachia* levels. Finally, *Wolbachia* levels were sensitive to changes in host transcription. We observed that disruption of components in the mediator complex and regulators of transcription from Polymerase II promoters led to decreased *Wolbachia,* whereas knockdown of the BRD4pTEFb complex involved in transcriptional pausing and other transcriptional elongation factors resulted in increased *Wolbachia* levels. Together, this manual curation revealed that the whole genome screen yielded host genes that suppress or enhance *Wolbachia* levels and that these primary hits could be classified into distinct gene categories.

**Figure 4.**
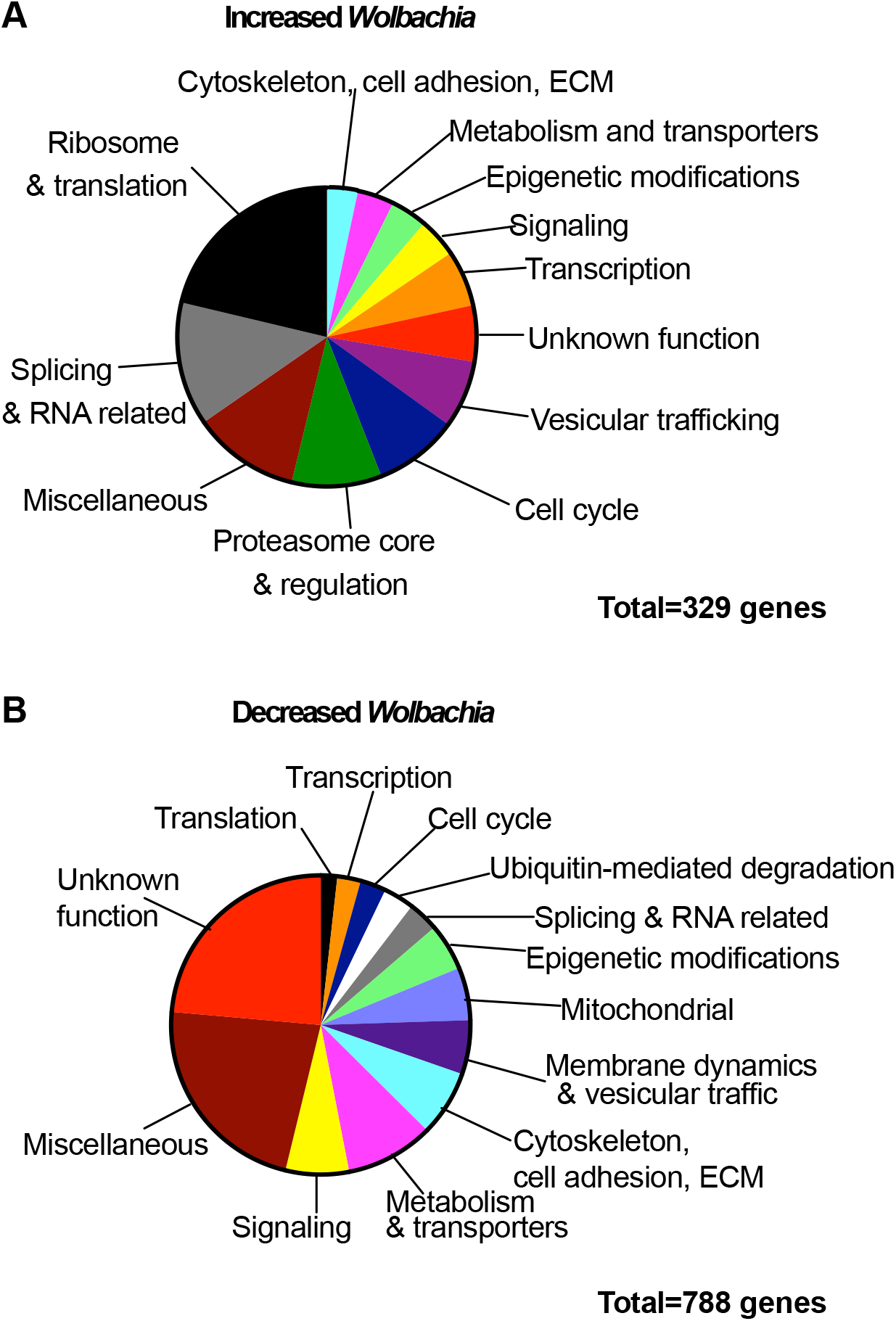
Primary screen hits classified into manually-curated gene categories and GO term analysis. **(A)** 329 primary gene hits that lead to increased *Wolbachia* levels upon RNAi knockdown manually curated into gene categories. **(B)** 788 primary gene hits that lead to decreased *Wolbachia* levels upon RNAi knockdown manually curated into gene categories.

Further GO term enrichment analysis using the online tool Panther™ (http://www.pantherdb.org/) suggested that the 329 genes resulting in *Wolbachia* increases formed a robust dataset as many of the enriched terms overlapped with our manual curation (S6 Fig). In contrast, there was a lack of enrichment for the 788 Wolbachia-decreasing genes even though manual curation had sorted many of these genes into categories. For this reason, further analysis focused on the 329 host genes that increased *Wolbachia* when knocked down by RNAi.

### Perturbations in host translation initiation and ribosome networks lead to increased *Wolbachia* levels

To assess whether specific host networks were enriched within the 329 host genes identified as potential suppressors of *Wolbachia* we used two bioinformatic tools namely the Kyoto Encyclopedia of Genes and Genomes (KEGG), and the protein complex enrichment analysis tool (COMPLEAT) with criteria for a network restricted to complexes with 3 or more components (p<0.05) [51]. This analysis revealed enrichment of several host networks among the 329 genes whose knockdown resulted in *Wolbachia* increases including a striking 67.5% of the core cytoplasmic ribosome (56/83 expressed subunits) and many translation initiation components (Fig 5A, and S7 Fig). These findings strongly suggested that perturbations in host translation components could alter *Wolbachia* levels. For both networks, the majority of components did not significantly affect cell growth within the duration of the RNAi screen assay (circles), though some did have a negative impact (robust Z<-1) (square) (Fig 5A, see Fig 3D). Importantly, these data show that *Wolbachia* level fluctuations are independent of host cell growth changes because *Wolbachia* levels increased in RNAi knockdowns of network components regardless of the presence or absence of cell growth changes (Fig 5A). We chose to validate and characterize the novel *Wolbachia-host* translation interaction identified in the whole genome RNAi screen.

**Figure 5.**
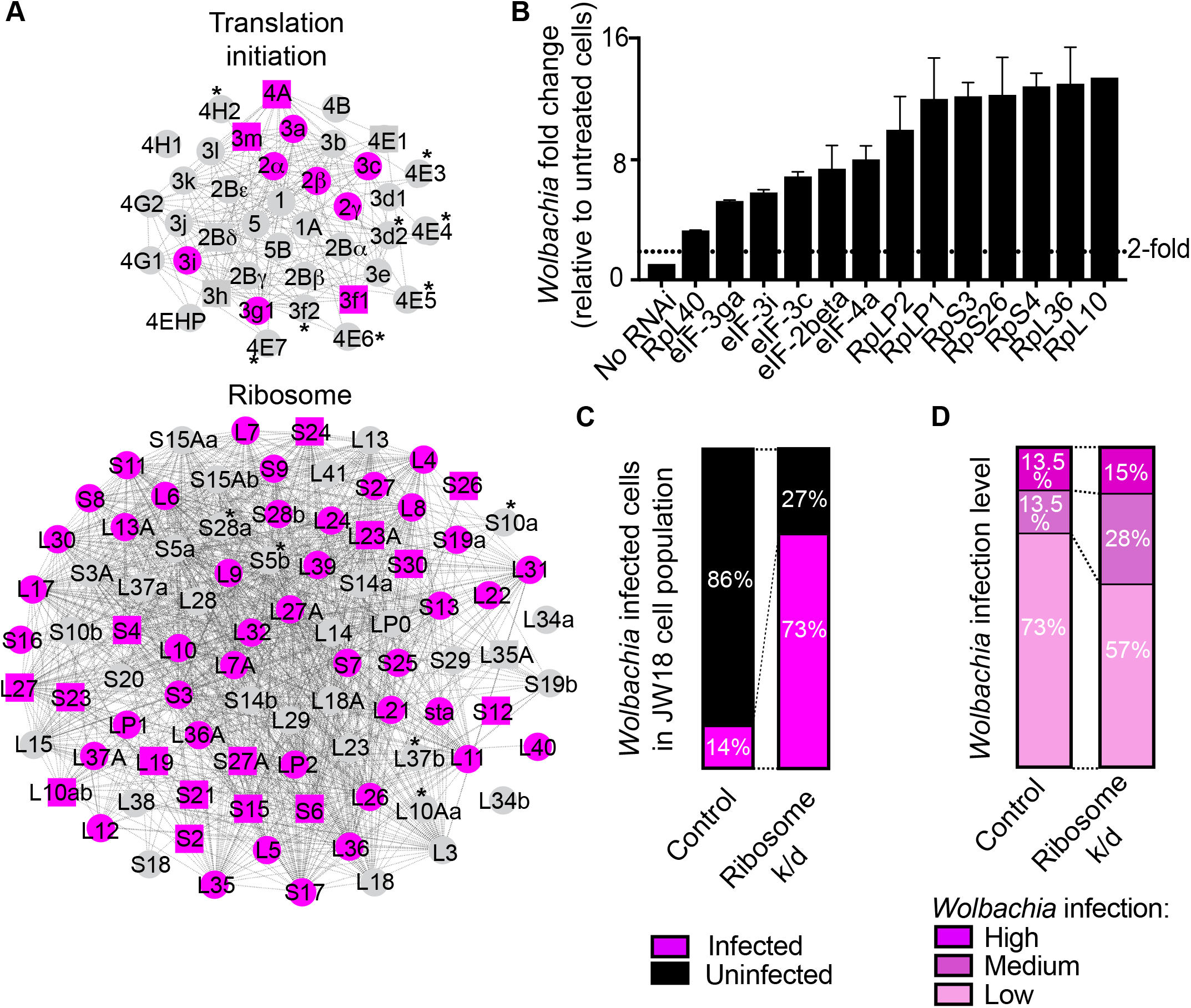
Host translation initiation and ribosome networks suppress *Wolbachia* levels. **(A)** The host translation initiation and ribosome networks identified by COMPLEAT and KEGG analysis are enriched for genes that increase *Wolbachia* levels when knocked down by RNAi in the primary screen. *Wolbachia* level changes are indicated by color: increases (magenta) and no effect (grey). Asterisks mark subunits that are not expressed in the JW18 cell line. Changes in cell growth in the whole genome screen assay are indicated by icon shape: no change (circle) or decrease (square). **(B)** Representative genes from each network were validated by RNAi. Effects on *Wolbachia* levels were assessed quantitatively by a DNA qPCR assay measuring the number of *Wolbachia* genomes (wspB copy number) relative to *Drosophila* nuclei (RpL11 copy number). Network validation is represented relative to untreated JW18 cells. **(C)** Quantification of *Wolbachia-infected* (magenta) and uninfected (black) cells within the JW18 cell population in control and ribosome *(RpS3)* RNAi knockdown conditions. *Wolbachia* infection was detected using the *Wolbachia-detecting 23s rRNA* FISH probe and cells were identified by DAPI staining of host nuclei. **(D)** Classification of the level of *Wolbachia* infection within infected cells of the JW18 cell population under control and ribosome (*RpS3*) knockdown conditions (seen in C). For each cell population >500 cells were quantified for *Wolbachia* infection level by the following criteria: low (1-10 *Wolbachia),* medium (11-30 *Wolbachia),* and high infection (>30 *Wolbachia).*

We validated the influence of the ribosome and translation initiation complex on *Wolbachia* levels by knocking down representative members of each network using RNAi knockdown in JW18 cells (Fig 5B). Each gene was validated using two different dsRNA amplicons that were designed to target different parts of the gene. Effects on *Wolbachia* levels were assessed quantitatively by DNA qPCR measuring the number of *Wolbachia* genomes *(wspB* DNA copies) relative to the number of host cell nuclei (RpL11 DNA copies). Network validation is represented relative to untreated JW18 control cells (No RNAi) and the positive control *RpL40* RNAi knockdown is included for reference. For the translation initiation network, we selected *eIF-4a, eIF-2 subunit beta, eIF-3c, eIF-3i,* and *eIF-3ga.* All subunits’ RNAi knockdown significantly increased *Wolbachia* levels by 5-fold or more (Fig 5B). For the ribosomal network, we selected *RpL10, RpL36, RpLP1, RpS4, RpLP2, RpS3* and *RpS26* for validation and each RNAi knockdown resulted in a significant increase of nearly 10-fold or higher *Wolbachia* levels relative to untreated JW18 control cells (Fig 5B). In control culturing conditions, JW18 cells maintain a stable *Wolbachia* infection level within the population when cells are split 1:1 every 4 days implying that the cells as well as the *Wolbachia* double in this timeframe. Thus, we would expect that if knockdown of a host gene results in slowed cell growth then the resulting increase in *Wolbachia* would be a two-fold increase at most. Instead, our results show far greater increases in *Wolbachia* levels. As such, we suggest that the *Wolbachia* level increases observed were mostly independent of host cell growth rate. To summarize, we were able to validate that RNAi knockdown of translation initiation and ribosomal networks leads to striking increases in *Wolbachia* levels in JW18 cells.

The increases in *Wolbachia* levels from the knockdown of these host networks could be a result of increased infection within cells that were already infected, increased cell-to-cell spreading, or a combination of both. To test these possibilities, we characterized the *Wolbachia* infection in the JW18 cell populations for ribosome RNAi knockdown (RpS3 subunit) compared to non-knockdown control JW18 cells. We visually classified the level of *Wolbachia* infection in cells within each population using the *Wolbachia-detecting 23s rRNA* FISH probe combined with DAPI staining and the *GFP-Jupiter* transgene labelling microtubules to identify the cells. Each cell was classified according to its *Wolbachia* infection into the following categories: uninfected (no *Wolbachia),* low (1-10 *Wolbachia),* medium (11-30 *Wolbachia),* and high (>30 *Wolbachia)* infection. Similar to Figure 1C, in a control no-knockdown JW18 control population 13% of the total number of cells were infected. Of the infected cells in the control population, 73% had a low level of infection whereas 13.5% had a medium level infection and 13.5% had a high level of infection (Fig 5C, D). In contrast to the control, RNAi knockdown of the ribosome RpS3 subunit resulted in an overall dramatic increase in the total number of infected cells (73%) within each population (Fig 5D). A comparison of the number of medium and highly infected cells revealed a 1.6-fold increase in the number of medium and highly infected cells in the network knockdown conditions compared to the *LacZ* knockdown control (Fig 5D). Our results show an increase in the number of medium and high infection cells within the ribosomal network knockdown cell populations, however the majority of infected cells contain a low level of infection suggesting that the major factor for the dramatic increased *Wolbachia* levels is most likely the result of increased cell-to-cell spreading.

Next, we tested whether these networks could influence *Wolbachia* in the fly (Fig 6 and S8 Fig). In *Drosophila, Wolbachia* are found abundantly in the ovary. To test the effect of perturbing the ribosome, females from a *Wolbachia-infected* stock were crossed to available ribosomal stocks carrying a mutant allele for RpL27A and RpS3 at 25°C.

**Figure 6.**
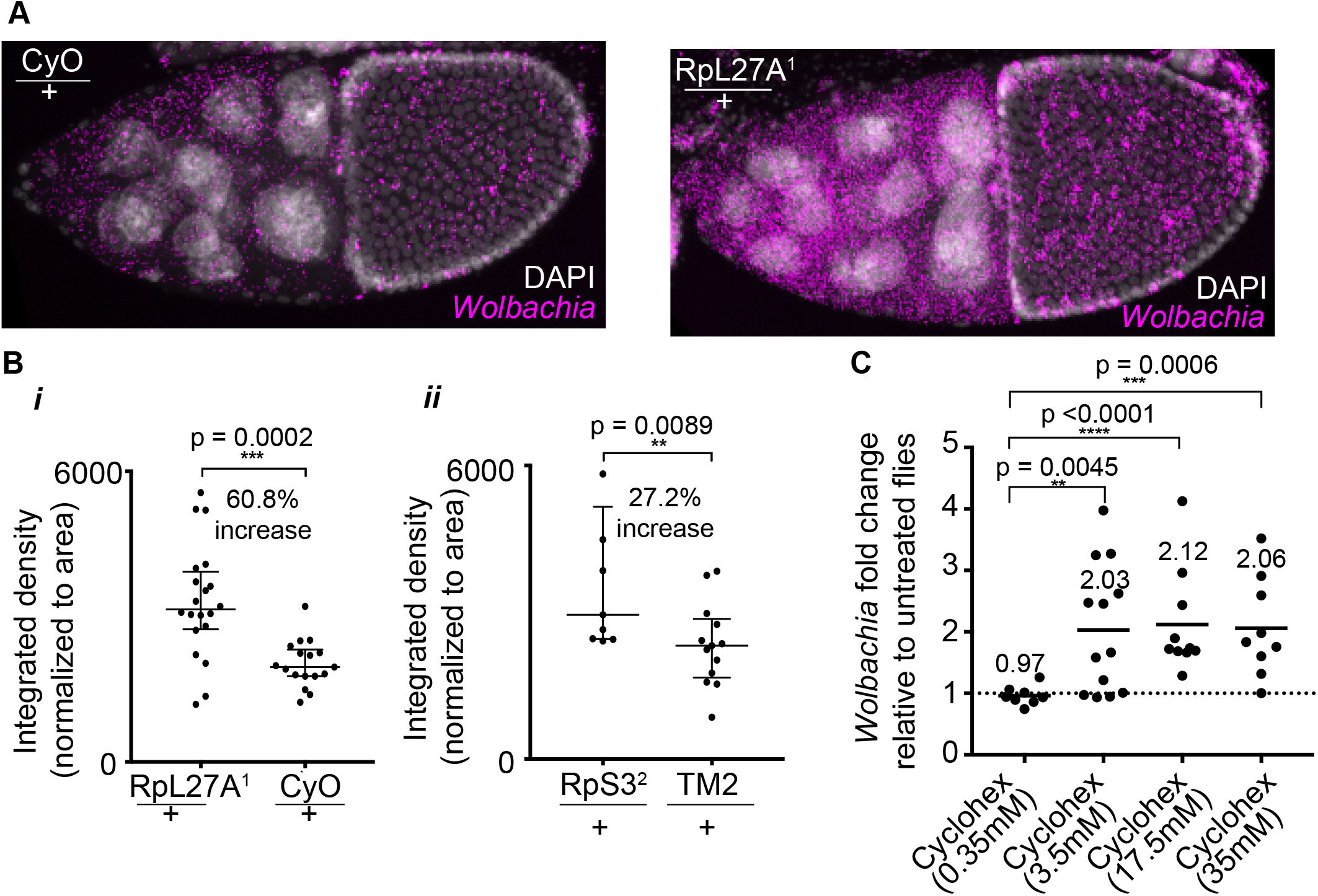
Perturbations in host ribosome or global translation lead to increased *Wolbachia* levels in the fly. **(A)** *Wolbachia-infected* stage 10 *Drosophila* egg chamber of control sibling (CyO/+) and ribosome mutant sibling (RpL27A^1^/+) processed to visualize *Wolbachia* using a *23s rRNA* FISH probe (magenta) and cell nuclei (DAPI). (B) Quantification of integrated density of the *Wolbachia* FISH probe in stage 10 egg chambers collected from 15-25 *Drosophila* ovary pairs for each genotype. Differences between control and mutant siblings is statistically significant (Mann Whitney, p<0.001). (C) *Wolbachia-infected Drosophila* flies fed on cycloheximide-containing food or control food for 7 days. Cycloheximide-fed flies displayed significantly increased *Wolbachia* levels in the whole fly as measured by DNA qPCR (Mann Whitney, p<0.001). Results displayed as fold-change relative to control food fed flies. Each data point represents an individual fly. Bars represent mean fold change.

Then, the *Wolbachia* infection level in the ovaries of 5 day-old *Wolbachia-infected* siblings were compared by RNA FISH for *Wolbachia 23s rRNA.* We observed dramatic increases in *Wolbachia* levels in the ribosomal mutants compared to the control sibling ovaries (Fig 6A). Quantification of the integrated density of the *23s rRNA Wolbachia* FISH probe in Z-stack projections of stage 10 egg chambers for the ribosomal mutants confirmed a 60.8% (RpL27A) and 27.2% (RpS3) increase compared to their respective control siblings (Fig 6B) (Non-parametric Mann Whitney, RpL27A p=0.0002, RpS3 p=0.0089). In conclusion, these results demonstrate that *Wolbachia* levels are sensitive to changes in the host ribosomal network in the *Drosophila* ovary.

Finally, we asked whether a direct relationship exists between *Wolbachia* and host translation. To do this we asked whether chemical inhibition of host translation by cycloheximide would alter *Wolbachia* levels in host *Drosophila. Wolbachia-infected D. melanogaster* were fed on cycloheximide-containing food or control food for 7 days prior to genomic DNA extraction of whole flies. We tested the *Wolbachia-levels* in individual whole flies using DNA qPCR and found increased *Wolbachia* levels in flies fed on cycloheximide compared to control flies (Fig 6C). This suggested that *Wolbachia* levels are sensitive to host translation and that perturbation of host translation leads to increased *Wolbachia* levels.

### Increased *Wolbachia* levels correlate with low host translation

Having observed that *Wolbachia* levels are sensitive to host translation, we wanted to observe the relationship between *Wolbachia* and host translation levels in an unperturbed manner in *Drosophila* JW18 cells and in the fly. To correlate levels of host translation with levels of *Wolbachia,* we combined *Wolbachia* RNA FISH detection with a visual fluorescent ‘click’ chemistry-based method to assess global protein synthesis levels in host cells (Fig 7). This assay is based on a sensitive, non-radioactive method that utilizes ‘click’ chemistry to detect nascent protein synthesis in cells (Fig 7A) [52]. Detection of protein synthesis was based on the incorporation of a specialized alkyne-modified methionine homopropargylglycine (HPG) or alkyne-modified puromycin (OPPuro) into newly synthesized proteins in JW18 cells or *Drosophila* testes respectively. Labelled proteins were detected using a chemo-selective ligation or “click” reaction between the alkyne modified proteins and an azide-containing fluorescent dye which was added. This resulted in a fluorescent readout within each host cell correlating to the level of protein synthesis. Note that we assumed the majority of protein synthesis detected in this assay was host-related, however HPG can also be incorporated during bacterial protein synthesis, thus *Wolbachia* translation will have contributed to the overall fluorescent readout. We further processed the samples to detect *Wolbachia* by RNA FISH and then imaged cells using confocal microscopy (Fig 7A, B). Quantification of this fluorescent readout of protein synthesis in the JW18 population revealed that cells containing *Wolbachia* consistently had lower levels of global translation (Fig 7C). This observation is consistent with previous observations of a global reduction in host translation [53] and translation machinery [49, [49, 53, 54] in the presence of *Wolbachia* infection. To extend this observation, we correlated the level of global host translation with *Wolbachia* infection level. We found that increasing levels of *Wolbachia* infection resulted in significantly lower levels of protein synthesis with the median translation level decreased by 43% comparing cells that did not contain *Wolbachia* to cells containing a medium-high level of infection (Fig 7C). Together, these results show a negative correlation between *Wolbachia* levels and host translation. Finally, we asked if this correlation was also observed in the fly. For this, we took advantage of the *Drosophila* testis hub located at the tip of the testis. The hub is the niche that maintains surrounding germline stem cells (GSCs) (Fig 7D) and translation is low in the hub compared to the attached germ cells, even in uninfected flies[55]. We assessed translation levels *in vivo* in infected *wMel Wolbachia Drosophila* males using click chemistry that incorporates a fluorescently-detected alkyne-labeled puromycin (OPPuro). We found *Wolbachia* concentrated in the testis hub (yellow outline) and outside sheath cells (grey outline) which had very low levels of translation compared to surrounding GSCs (Fig 7E, F). This observation provides an *in vivo* example where high *Wolbachia* levels correlate with low levels of host translation. Together our results suggest a consistent inverse correlation between *Wolbachia* levels and host translation, whereby low translation in the host favors high Wolbachia density and Wolbachia levels may influence host translation.

**Figure 7.**
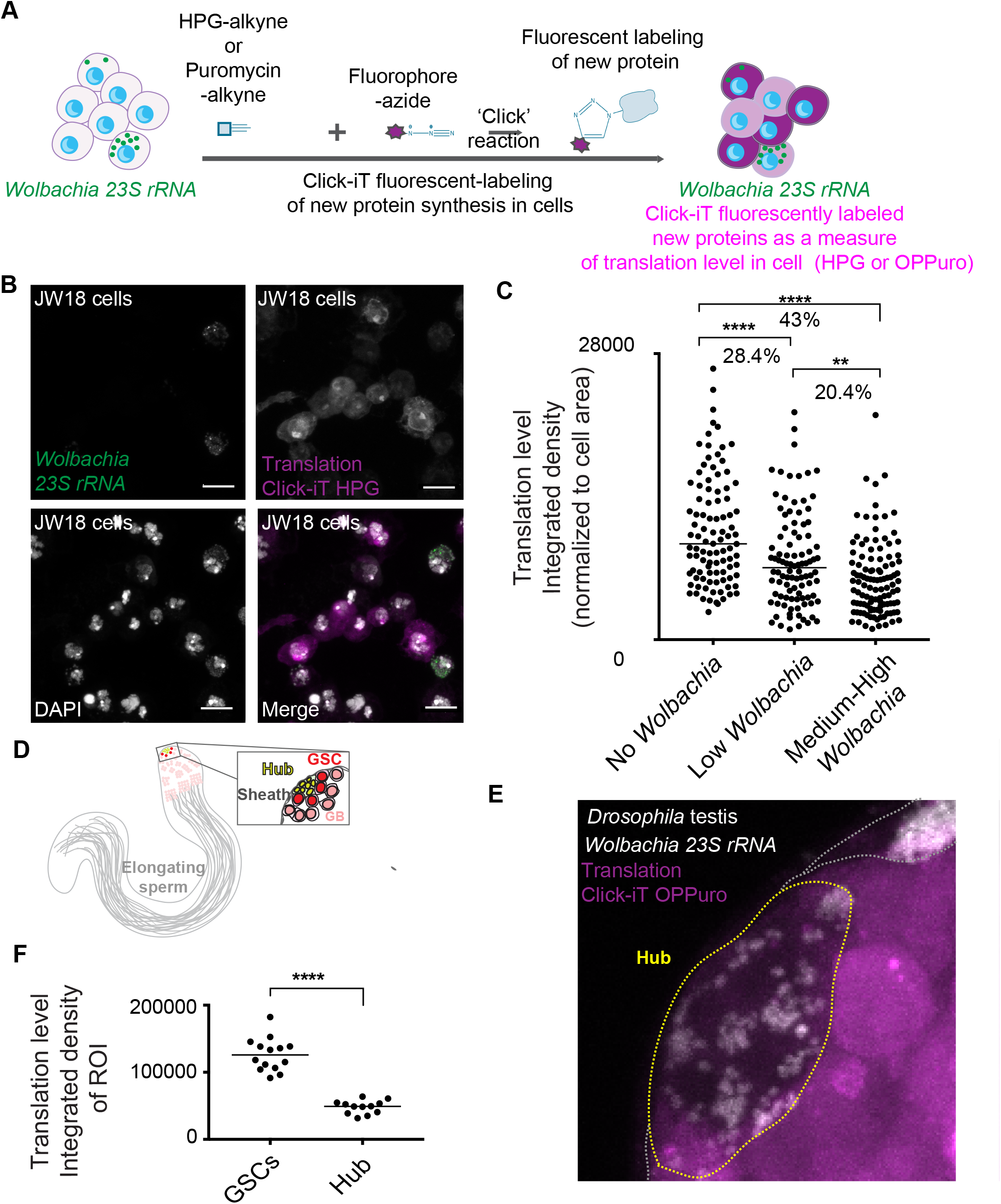
*Wolbachia* levels negatively correlate with host translation levels. **(A)** Illustration of Click-iT method that fluorescently labels newly synthesized proteins in host cells as a measure of translation level. JW18 cells were incubated with a modified methionine called HPG containing an alkyne or whole testes were incubated with a modified puromycin containing an alkyne. In each case these alkyne-containing reagents were incorporated into newly synthesized proteins during incubation. Thereafter, samples were processed for the ‘click’ reaction by adding an azide-tagged fluorophore (purple) which reacted with the incorporated alkynes to label new proteins fluorescently (purple). The fluorescent intensity is used as a measure of translation level within the host cell. This assay is combined with *Wolbachia-detecting 23s rRNA* FISH assay (green). **(B)** JW18 cells labeled by Click-iT HPG (purple), DAPI (white), and *Wolbachia-detecting 23s rRNA* FISH assay (green). **(C)** Quantification of HPG Click-iT assay fluorescent signal for protein synthesis within individual cells of the JW18 population. Each dot represents a single cell. Cells were imaged at 63x to capture the whole cell using Z-stack imaging. The integrated density of the fluorescent signal for each cell was calculated using Fiji and signal was normalized for the size of each cell. We categorized cells according to *Wolbachia* infection level: no *Wolbachia,* low *Wolbachia* (1-10 bacteria), or medium-high *Wolbachia* (11 bacteria or more). Translation level is significantly decreased in cells that contain *Wolbachia.* The higher the level of *Wolbachia,* the lower the translation level. (p<0.001). **(D)** The *Drosophila* testis illustration highlighting the stem cell niche (hub) at the tip of the testes surrounded by and directly contacting the germline stem cells (GSCs) (dark red). GSCs divide perpendicular to the hub to maintain one GSC that touches the hub and one daughter cell (light red) that matures into a developing sperm. The testes is surrounded by sheath cells (grey). **(E)** Projected Z-stack of a *Wolbachia-infected Drosophila* testis hub (yellow outline). Testes were treated with Click-iT OPPuro to fluorescently label newly synthesized proteins (purple) and *Wolbachia* are labeled by FISH (white). *Wolbachia* mainly occupy the hub niche and sheath cells (grey outline) which have low levels of translation as measured by Click-iT OPPuro assay (purple). **(F)** Quantification of HPG OPPuro assay fluorescent signal for protein synthesis measured in ImageJ as integrated density in the hub compared to surrounding GSCs.

## Discussion

The recent applications of *Wolbachia* as a tool to lower the transmission of vector-borne viruses necessitates a comprehensive analysis of the relationship between *Wolbachia* and the vector host. In particular, the observation that increasing *Wolbachia* density leads to stronger antiviral effects in vectors [10, 25–27] argues for a thorough examination of how intracellular *Wolbachia* levels are controlled. Here we focused on understanding which host systems influence *Wolbachia* levels. We performed a comprehensive unbiased whole genome RNAi screen that adapted RNA FISH for a high throughput approach. Traditionally, visual cell culture-based screens that investigate host-pathogen interactions use immunofluorescent staining, luminescent readouts, or fluorescently-tagged pathogens. The lack of tools for *Wolbachia* such as a commercially available antibody or a fluorescently-tagged *Wolbachia* strain necessitated our RNA FISH approach as a visual assay. This screen confirmed the feasibility of an RNA FISH detection approach as 1117 host genes were identified that alter *Wolbachia* levels. This accounted for approximately 8% of all screened genes. Knock down of 329 of these genes resulted in increased *Wolbachia* levels whereas 788 genes resulted in decreased *Wolbachia* levels. In summary, the screen successfully identified a comprehensive array of host genes that influence intracellular *Wolbachia* levels.

Here we report that *Wolbachia* levels are sensitive to changes in host translation. When host translation components such as the ribosome or translation initiation complex are perturbed by RNAi we observe remarkable increases in *Wolbachia* levels (Fig 5). In support, *Wolbachia* levels increase in the *Drosophila* ovary of ribosomal mutants and *Wolbachia* levels increase upon global host translation inhibition when flies were fed with cycloheximide (Fig 5, 6). Collectively, these results provide the first evidence that *Wolbachia* levels are sensitive to host translation level changes and suggests that host translation might normally play an inhibitory role in regulating intracellular *Wolbachia* levels.

In addition to the sensitivity that *Wolbachia* displays towards host translation levels it is possible that host translation is also directly affected by *Wolbachia.* Quantification of protein synthesis in individual JW18 cells revealed that host cells containing *Wolbachia* had dampened host translation (Fig 7). When classified according to *Wolbachia* infection level, higher *Wolbachia* levels correlated with significantly lower levels of host translation as measured by global protein synthesis levels. We did not observe changes in translation components at the RNA level (S9 Fig), however, recent proteomics studies revealed that over 100 host proteins with roles in host translation were suppressed in the presence of *Wolbachia* [49, 56]. The mechanism/s underlying this remains to be determined. One possibility is that the host translation is dampened by a stress response to *Wolbachia.* However, our gene expression analysis did not suggest any major alterations in stress response-related genes at the RNA level in response to *Wolbachia* (S10 Fig). Yet, several significant changes in stress response were detected at the proteome level suggesting that stress could play a role in the *Wolbachia-host* intracellular relationship [49, 56, 57]. Host translation shutdown via metabolic stress pathways is a common mechanism employed by pathogens [58]. The other possible mechanism for dampening host translation is active manipulation of host translation machinery by *Wolbachia* perhaps at the post-translational level as our data do not suggest changes at the transcriptional level (S9 Fig). *Wolbachia* encodes and expresses a fully functional type IV secretion system and many potential effector proteins [59–61]. Although the majority of *Wolbachia* effector proteins remains to be characterized, it is possible that *Wolbachia* encodes effector/s that can manipulate the host’s translation machinery at a post-translational level as is the case for other intracellular bacteria such as *Legionella* [62, 63].

*Wolbachia* interaction with host translation could be important in the context of positive-strand RNA virus infection in the host. *Wolbachia-mediated* suppression of viral replication in hosts is well described [7, 8, 11, 12, 64]. Multiple mechanisms may underlie this observation including interference with viral entry and very early stages of viral replication [38, 53, 57, 64–71]. All viruses depend on host translation machinery for replication of their genomes. One intriguing possibility is that the interaction between *Wolbachia* and host translation could impact viral replication [53]. *Wolbachia* infection inhibits positive-strand RNA viral replication at very early stages of viral replication in the virus lifecycle [53, 64]. Thus, changes in host translation could be one mechanism by which *Wolbachia* infection contributes towards viral replication interference. Future work to elucidate whether this is a contributing mechanism to the *Wolbachia-mediated* antiviral response in a wide range of Wolbachia-host-virus relationships may provide valuable field applications for combating vector-borne viruses.

Our whole genome screen yielded a diverse range of host systems and complexes that influenced *Wolbachia* levels. Manual curation and bioinformatic analyses such as GO term enrichment and network analysis identified host pathways such as translation initiation, ribosome, cell cycle, splicing, immune-related genes, proteasome complex, COPI vesicle coat, polarity proteins and the Brahma complex. The GO term enrichment analysis and COMPLEAT network analysis suggested that the 329 genes resulting in *Wolbachia* increases formed a more robust dataset than the larger 788 gene category resulting in *Wolbachia* decreases owing to a lack of enrichment for specific networks and processes in this category. For this reason, we focused on the host networks that increased *Wolbachia* in this report. Nevertheless, future follow-up analysis on genes that decreased *Wolbachia* levels especially the larger categories such as metabolism & transporters, cytoskeleton, cell adhesion & extracellular matrix, as well as membrane dynamics and vesicular trafficking may yield rewarding results. We already appreciate that *Wolbachia* relies on several aspects of these broad categories. For example,

*Wolbachia* can alter host iron, carbohydrate and lipid metabolism [72–76]. Further, *Wolbachia* interacts with host cytoskeleton such as microtubules for transport and host actin [32, 34, 35, 77]. Finally, *Wolbachia* resides within a host-derived membrane niche, as such genes identified in the membrane dynamics and vesicular trafficking category would be of interest [78, 79]. Further investigation of these Wolbachia-decreasing categories may provide comprehensive insights into *Wolbachia-host* interactions.

Interestingly our study also revealed that knockdown of the core proteasome leads to increases in *Wolbachia* levels (Fig 4A, S7 Fig, S8 Fig). A previous report suggested that *Wolbachia* require high levels of proteolysis for optimal survival [36]. An explanation for this discrepancy might be that previous observations of decreased *Wolbachia* levels were based on RNAi experiments that knocked down ubiquitin-related components not the core proteasome. Ubiquitination is known to function in many diverse contexts and pathways such as autophagy, cell cycle, immune response, DNA damage response and regulation of endocytic machinery [80]. Our screen also identified several ubiquitin-related components whose knockdown resulted in decreased *Wolbachia* levels (Fig 4B). Perhaps *Wolbachia* relies on the host ubiquitination system for survival in an unknown but specific context, not simply for providing amino acids as nutrients from the degradation of proteins by the proteasome. Our gene expression data (S9 Fig) along with recent proteomics studies [49, 56] suggest that the host proteasome is upregulated in the presence of *Wolbachia.* We propose that the host proteasome plays an inhibitory role in *Wolbachia* level regulation. Perhaps *Wolbachia* levels are controlled by degradation of effector proteins in the cytosol, thereby preventing *Wolbachia* from utilizing the host cell in an optimal manner. The results both in the JW18 cell line as well as in the ovary of *D. melanogaster* strongly suggest that the host core proteasome normally plays a restrictive role in *Wolbachia-host* interactions that is separate from observations of ubiquitin pathway perturbation.

In summary, here we presented a whole genome screen to identify host systems that influence *Wolbachia* levels. Our focus was on *Wolbachia* sensitivity to alterations of host translation-related components such as the ribosome and translation initiation factors. We report a novel relationship between *Wolbachia* and host translation and suggest a restrictive role for host translation on *Wolbachia* levels. Future work to identify whether *Wolbachia* is able to actively manipulate host translation will provide valuable insight into understanding this unique host-symbiont relationship.

## Materials and Methods

### Cell culture

A stable *Wolbachia-infected Drosophila* cell line (JW18) and a tetracycline-treated *Wolbachia-free* cell line (JW18TET) (kindly provided by William Sullivan at UCSC) were maintained in Sang and Shield media (Sigma) supplemented with 10% heat inactivated One Shot™ Fetal Bovine Serum (Life Technologies).

### Fly stocks and husbandry

*D. melanogaster* infected with the *wMel Wolbachia* strain was a gift from Luis Teixeira (Instituto Gulbenkian de Ciência). A *Wolbachia-free* version of the line was generated by raising two generations on food containing 0.05 mg/ml of tetracycline hydrochloride (Sigma). Virgin adult females were collected and individually crossed with males to establish isofemale lines. Lines were not used in experiments for at least one year following tetracycline-treatment. To generate a *Wolbachia-infected* double balancer line, *Wolbachia-infected* virgins were crossed to Sp/CyO; MKRS/TM6b males. In the next generation *Wolbachia-infected* +/CyO;+/TM6b female virgins were crossed to males of the original double balancer stock. In the final generation, a stock of *Wolbachia-infected* Sp/CyO; MKRS/Tm6b double balancers was established.

Ribosomal mutant fly stocks were ordered from the Bloomington *Drosophila* Stock Center. The following haploinsufficient lines were used: RpS3^2^/TM2 (stock no. 1696) and RpL27A^1^/CyO (stock no. 5697). Males from each line were crossed at 25°C to the *Wolbachia-infected* double balancer line described above. Siblings from each cross were matured for 5 days before ovaries were dissected and stained for *Wolbachia* using the 23s rRNA *Wolbachia-specific* FISH probe. Stage 10 egg chambers were imaged in Z-stacks using confocal microscopy. Quantification of the integrated density of the *23s rRNA Wolbachia* FISH probe in stage 10 egg chamber Z stacks were done using Fiji Image Processing software as described in the FISH section.

A dominant temperature-sensitive (DTS) lethal mutant for proteasome subunit *Pros26,* known as *DTS5,* was a gift from John Belote, Syracuse University. Heterozygotes die as pupae when raised at 29°C, but are viable and fertile at 25°C. The mutant contains a missense mutation in the gene encoding the β6 subunit of the 20S proteasome. Males were crossed to *Wolbachia-infected* female double balancers at the permissive temperature of 25°C. Hatched offspring from the cross were matured at the non-permissive 29°C for 5 days prior to dissection of the ovaries. Imaging and analysis of stage 10 egg chambers of siblings were done as described above for the ribosomal mutant crosses.

### Genomic DNA extraction and DNA qPCR for *Wolbachia* level quantification

Genomic DNA was extracted from cells or *Drosophila* tissues using a DNeasy Blood & Tissue Kit (Qiagen) following manufacturer’s instructions. To quantify the level of *Wolbachia* in the sample a DNA qPCR assay was performed using SYBR Green I Master 2x (Roche), using a Roche LightCycler 480 machine. Primer sets included a primer set to detect *wspB* which is a gene encoding a *Wolbachia* surface antigen (F: 5’ ACA ACA GCT ATA GGG CTG AAT TGG AA 3’, R: 5’ TCA GGA TCC TCA CCA GTC TCC TTT AG 3’), as well as a primer set to detect the *Drosophila* gene *RpL32* (also known as *RpL49)* (F: 5’ CGA GGG ATA CCT GTG AGC AGC TT 3’, R: 5’GTC ACT TCT TGT GCT GCC ATC GT 3’). *Wolbachia* levels were normalized by the host nuclear marker for each sample.

### RNA extraction, Reverse Transcription and RT-qPCR using JW18 cells

RNA was extracted and DNase-treated using a RNeasy kit (Qiagen) according to manufacturer’s instructions. Total RNA was reverse transcribed using an RNA to cDNA EcoDry™ Premix (OligodT) or EcoDry™ Premix (Random hexamer) (Clontech) according to manufacturer’s instructions. Quantitative PCR was performed on 1/200 of the RT reaction using LightCycler 480 SYBR Green I Master 2x (Roche) and a Roche LightCycler 480 machine. Results were normalized to the housekeeping gene *Rp49.* Primer sets used to validate RNAi knockdown were designed to amplify areas outside of the dsRNA amplicon. Gene knockdown was represented relative to expression levels in *LacZ* dsRNA-treated cells.

### DNA and RNA sequencing

Genomic DNA was extracted from JW18 cells in duplicate. Samples were quantified using a Qubit fluorometer (Thermo Fisher Scientific). DNA libraries were prepared using a Nextera DNA Library Prep kit (Illumina) according to manufacturer’s instructions. DNA libraries were sequenced on an Illumina HiSeq2500 Sequencing platform in two lanes as paired-end reads 100 cycle lanes.

Total RNA was extracted in triplicate from JW18 and JW18TET cells and DNase-treated. RNA was quantified by Nanodrop and 5μg of each sample was subjected to two rounds of rRNA depletion using a Ribo-Zero rRNA Removal Magnetic kit (Epicentre, Illumina) or NEBNext rRNA Depletion Kit (Human/Mouse/Rat) (New England BioLabs, E6310L). After rRNA depletion libraries were prepared according to manufacturer’s instructions using the NEBNext Ultra Directional RNA Library Prep Kit for Illumina (New England BioLabs, E7420L) and NEBNext Multiplex Oligos for Illumina Index Primers Set I (Illumina, E7335). After adaptor ligation, the libraries were amplified by qPCR using the KAPA Real-time amplification kit (KAPA Biosystems). Finally, libraries were purified using Agencourt AMPure XP beads (Beckman Coulter) as described in the NEBNext Ultra Directional RNA Library Prep Kit for Illumina (New England BioLabs, E7420L) protocol. Quality and quantity was assessed using a Bioanalyzer (Agilent) and a Qubit fluorometer (Thermo Fisher Scientific). Libraries were sequenced on an Illumina HiSeq2500 Sequencing platform in single read 50 cycle lanes.

### RNA-seq analysis

Differential gene expression analysis was performed from one lane of high output, single end reads 50, Illumina HiSeq run. The experiment consisted of 3 replicates each for JW18 and JW18TET cells. The alignment program, Tophat (version 2.0.9) (https://ccb.jhu.edu/software/tophat/index.shtml) was used for reads mapping with two mismatches allowed. Featurecounts (http://bioinf.wehi.edu.au/featureCounts/) was used to find the read counts for annotated genomic features. For the differential gene statistical analysis, DESeq2 R/Bioconductor package in the R statistical programming environment was used (http://www.bioconductor.org/packages/release/bioc/html/DESeq2.html).

### Copy number analysis of JW18 cell line and *Wolbachia*

We mapped short reads generated from DNA-Seq with Bowtie2 version 2.2.9 [81]. We used default parameters and mapped to combined sequences of *Drosophila* genome release 6 [82] and *Wolbachia pipientis wMel* ([59], GenBank accession ID AE017196.1). We determined basal ploidy level of JW18 cells by clustering normalized DNA-Seq read densities as in [83]. In doing that, we identified different copy number segments whose normalized read densities are between zero (no DNA content) to the mean density (basal ploidy level). Clusters of such read densities indicate the minimum ploidy. From the determined basal ploidy, we called copy numbers of JW18 cell line genome using Control-FREEC version 5.7 [84] at 1 kb levels. We called copy numbers in an identical way to [83] but with this exception; we performed calling twice and combined the results. Control-FREEC performs GC contents-based normalization of DNA-Seq reads. Therefore, we set the minimum expected GC contents to be 0.30 for robust copy number calling of the cell line genome first. Then we underwent our analysis again with the minimum expected GC contents of 0.25 to increase sensitivity against the bacterial genome. In our reports, we combined copy number calls from the former for JW18 cells, and from the latter for *Wolbachia.* In S1 Figure, we used DNA-Seq results from [83] to call copy number calls on S2R+ and Kc167 cells. We re-analyzed the original data after mapping to the release 6 genome as above.

### Genomic analysis of JW18 cell line *Wolbachia* strain

The method described in [60] was used to analyze the genotype of the *Wolbachia* strain in the JW18 cell strain. Briefly, fastq sequences were mapped against a “holo-genome” consisting of the Release 5 version of the *D. melanogaster* genome (Ensembl Genomes Release 24, Drosophila_melanogaster.BDGP5.24.dna.toplevel.fa) and the *Wolbachia wMel* reference genome (Ensembl Genomes Release 24, Wolbachia_ endosymbiont_of_drosophila_melanogaster.GCA_000008025.1.24) [85, 86]. Holo-genome reference mapping was performed using bwa mem v0.7.5a with default parameters in paired-end mode. Mapped reads for all runs from the same sample were merged, sorted and converted to BAM format using samtools v0.1.19 [87]. BAM files were then used to create BCF and fastq consensus sequence files using samtools mpileup v0.1.19 (options −d 100000). Fastq consensus sequence files were converted to fasta using seqtk v1.0-r76-dirty and concatenated with consensus sequences of *Wolbachia-* type strains from [27]. Maximum-likelihood phylogenetic analysis on resulting multiple alignments was performed using raxmlHPC-PTHREADS v8.1.16 (options −T 12 −f a −x 12345 −p 12345 −N 100 −m GTRGAMMA) [88]. Copy number variants were detected by visual inspection of read depth across the *wMel* genome.

### Gene set enrichment

GO enrichment analysis were performed using PANTHER™ Version 12.0 (release 2017-07-10) (http://www.pantherdb.org/). The entire set of screened genes was used as the experimental background. Protein complex enrichment analysis was performed using COMPLEAT (http://www/flyrnai.org/compleat/). As the experimental background we used the entire set of screened genes. Complex size was limited to ≥3, with a p value filter of p<0.05.

### dsRNA synthesis

Outdated amplicons from the *Drosophila* RNAi Screening Center (DRSC) whole genome library 2.0 were identified using Updated Targets of RNAi Reagents (UP-TORR) (http://www.flyrnai.org/up-torr). For amplicons that could be transferred to Release 6, we followed the dsRNA *in vitro* synthesis protocol as described by the DRSC. The DNA templates were generated by PCR on genomic DNA extracted from the JW18 cells, genomic DNA from wild-type flies or pBlueScript SK (+) plasmid DNA (in the case of *LacZ*). All gene specific primer sequences were selected by the DRSC and the T7 promoter sequence (TAATACGACTCACTATAGGG) was added to the 5’ ends of all primer pairs. Gradient PCR reactions were performed with Choice Taq Mastermix (Denville Scientific Inc.) using 5ng of genomic DNA, 0.1ng of plasmid DNA, or 1:3 diluted PCR template DNA. PCR products were verified by electrophoresis on a 0.7% (w/v) agarose gel with the 1 kb PLUS ladder (Invitrogen) and only products with a clear single band were selected for IVT. IVT was performed according to manufacturer’s instructions for the MEGAscript T7 Transcription Kit (Ambion) using 8μl of amplified T7-flanked PCR product per reaction. dsRNA products were DNase-treated using Turbo DNase (Ambion) and purified with Qiagen RNeasy Mini spin columns (Qiagen) according to manufacturer’s protocols. Quality of purified dsRNA was assessed by electrophoresis on a 0.7% agarose gel, and concentration was determined by Nanodrop (Thermo Fisher Scientific) [40].

### RNAi in JW18 cell line

RNAi in JW18 cells was done using a bathing method described by the DRSC [40]. dsRNA aliquots were prepared in serum-free Sang and Shield media (Sigma). dsRNA was added to wells to yield a final concentration of 25nM. For the whole genome screen the pre-arrayed DRSC *Drosophila* Whole Genome Library Version 2.0 was used. For each RNAi experiment, sub-confluent JW18 cells were scraped, pelleted (1000rpm for 5-15 minutes), and re-suspended in serum-free Sang and Shield media (Sigma) to seed 40 000 cells in 384 well format. Cells and dsRNA were incubated together at room temperature for 30 minutes in serum-free conditions. Thereafter Sang and Shield media (Sigma) supplemented with 10% heat inactivated One Shot™ Fetal Bovine Serum (Life Technologies) was added to each well and incubated at 25°C for 5 days before analysis.

### Fluorescent *in situ* hybridization (FISH)

Cells were plated in Poly-L-lysine-coated chambered cover-glass wells (Thermo Scientific) and allowed to settle. Medium was aspirated and cells were washed with 1xPZBS before fixing with 4% PFA in 1xPBS (Electron Microscopy Sciences). Cells were washed twice in 1xPBS followed by two washes in 100% methanol (Fisher) before finally adding 100% methanol to each chamber and sealing it with Parafilm^®^ M Film (Sigma) for storage at −20°C overnight or up to 1 month. Samples were rehydrated using the following washes: MeOH: PBT (1xPBS, 0.1% Tween-20) (3:1), MeOH:PBS (1:1), MeOH: PBS (1:3), and a final wash in 1xPBS. Samples were then post-fixed for 10 minutes in 4% PFA at room temperature. In a pre-hybridization step, samples were incubated in 10% deionized formamide and 2x SCC for 10 minutes at room temperature. Pre-hybridization buffer was then removed, and a hybridization solution containing a *Wolbachia-specific* FISH probe was added and incubated overnight at 37°C. For each sample, the volume of hybridization buffer added was dependent on the type of well used, but enough should be added to cover the sample. Typically, 60μl of hybridization buffer comprised 10% Hi-Di^TM^ deionized formamide (Applied Biosystems Life Technologies), 1 μl of competitor (5mg ml^−1^ *E. coli* tRNA (Sigma) and 5 mg ml^−1^ salmon sperm ssDNA (Ambion)), 10mM vanadyl ribonucleoside complex (New England Biolabs), 2xSSC (Ambion), 50μg nuclease-free BSA (Sigma), 10ng *Wolbachia-specific* FISH probe, made up to 60μl with DEPC-treated water. The *Wolbachia* specific probe was designed to the *Wolbachia 23s rRNA* and labeled with Quasar670 (Stellaris). After overnight hybridization samples were washed twice in pre-warmed pre-hybridization buffer for 15 minutes at 37°C. Followed by two washes in 1x PBS for 30 minutes each. Finally, samples were stained for 5 minutes in 1:500 DAPI:1xPBS followed by two washes in 1x PBS. Unless otherwise stated, samples were imaged as Z-stacks on a Zeiss LSM 780 confocal at 63x. For *Wolbachia* detection in dissected *Drosophila* ovaries and testes, the same protocol was followed from fixation onwards.

Quantification of *Wolbachia* levels based on the intensity of the *Wolbachia 23s rRNA* probe for stage 10 egg chambers was done in Fiji Image processing software. Z-stacks capturing entire stage 10 egg chambers were projected as ‘sum slices’. Each stage 10 egg chamber was manually outlined using the Freehand tool. The measurements tool was set to capture the integrated density within the outlined egg chamber of the FISH probe channel stack as well as provide an area measurement of the outlined egg chamber. We normalized the integrated density reading for each by its total area.

### Automated whole genome RNAi screening in JW18 cells

Large-scale RNAi screening was done using the DRSC *Drosophila* Whole Genome Library Version 2.0 that was seeded in Corning clear bottom, black 384 well plates with 0.25μg dsRNA pre-arrayed per well. This concentration of dsRNA was appropriate for the bathing method of RNAi [40]. JW18 cells were re-suspended in serum-free media at 4×10^6^ cells/ml and an automated Matrix Wellmate dispenser (Thermo Fisher Scientific) was used to dispense 40 000 cells into each well of the 384 well plates in a sterile tissue culture hood. The cells were incubated with the dsRNA in serum-free media for 30 minutes before automatic dispensing of Sang and Shield media (Sigma) supplemented with 10% heat inactivated One Shot™ Fetal Bovine Serum (Life Technologies) into each well. Plates were incubated at 25°C for 5 days in a humidity chamber. After 5 days plates were drained followed by automated dispensing of 4% paraformaldehyde (Electron Microscopy Sciences) and incubation at room temperature for 10 minutes and automatically aspirated thereafter. An automated BioTek EL406 liquid handler (BioTek) was used throughout the protocol for all aspiration steps and a Matrix Wellmate (Thermo Fisher Scientific) was used for all dispensing steps. Next, plates were washed once with 1xPBS followed by three washes with 100% methanol (Fisher), sealed with Parafilm^®^ M film, and stored overnight at −20°C (or up to 1 month). All subsequent rehydration, postfixation, pre-hybridization and hybridization steps of the RNA FISH protocol described above were carried out in an automated manner. After overnight hybridization at 37°C the plates were washed twice with pre-warmed pre-hybridization buffer followed by incubation at room temperature for 30 minutes with 1xPBS/DAPI. Finally, plates were washed once with 1xPBS and 40μl 1xPBS was dispensed into all wells and plates were sealed with aluminum foil and stored at 4°C.

Plates were imaged with a 20x objective lens using an Arrayscan^TM^VTI Microscope (Cellomics) coupled with the automated image analysis software HCS Studio Cellomics Scan Version 6.6.0 (Thermo Fisher Scientific). Image acquisition involved identification of DAPI stained cell nuclei as primary objects, followed by application of a ring mask around the primary objects to identify *Wolbachia* associated with each cell as secondary objects. Segmentation of the objects was optimized to exclude any areas containing cell clumps. For each well, 1500 primary objects (DAPI cell nuclei) were acquired. RNAi screen primary data analysis and criteria for hit selection is summarized in S4 Figure.

### Click-iT Protein Synthesis analysis

Protein synthesis levels in JW18 cells was detected using a Click-iT^®^ HPG Alexa Fluor^®^ 594 Protein Synthesis Assay Kit (Molecular probes, C10429). Regular Sang and Shield media (containing methionine) (Sigma) was removed from JW18 cells. Cells were washed once in 1xPBS. A working solution of Click-iT^®^ HPG was prepared according to manufacturer’s instructions using methionine-free Grace’s Insect Medium (Thermo Scientific). Cells were incubated for 30 minutes in 50μM Click-iT^®^ HPG working solution. After incubation, cells were washed once in 1xPBS followed by fixation in 5% formaldehyde. To combine the protocol with RNA FISH detection of *Wolbachia* in the cells, we next proceeded to wash the cells twice in methanol followed by storing the sample in 100% methanol at −20°C overnight. From this point, we followed the RNA FISH protocol described in the FISH section. After FISH hybridization and post-hybridization washes, we incubated cells with 0.5% Triton^®^ X-100 in 1xPBS for 20 minutes at room temperature. Cells were washed twice with 3% BSA in 1x PBS. The Click-iT^®^ Reaction Cocktail was added to the samples for 30 minutes at room temperature protected from light. Thereafter, samples were washed once with Click-iT^®^ Reaction Rinse Buffer before staining with 1x HCS NuclearMask™ Blue Stain working solution as per manufacturer’s instructions. Samples were imaged on confocal at 63x magnification. Controls included in the assay were as follows: incubation of cells with cycloheximide at a final concentration of 100μg/ml for 1 hour prior to the start of the experiment as well as during the 30-minute incubation with HPG; and a negative control sample that was not incubated with HPG.

Quantification of the Click-iT fluorescent intensity signal within each cell was done is a similar manner as described for FISH signal in egg chambers in the FISH section. Briefly, projected Z-stacks were manually outlined in Fiji and integrated density and area measurements were captured for each cell using the measurements tool. This allowed for a normalized integrated density measurement for individual cells. These data could then be paired with the *Wolbachia* level within individual cells as measured by RNA FISH.

Protein synthesis in the *Drosophila* testis was detected by the Click-iT Plus OPP Alexa Fluor 594 protein synthesis assay kit (Molecular Probes) as previously described [55]. Samples were incubated for 30 minutes in 1:400 Click-iT OPP reagent in fresh Shields and Sang M3 Insect medium (Sigma).

## Acknowledgements

We are grateful to Bill Sullivan (UCSC) for sharing the JW18 cell line. We thank Colin D. Malone for fruitful early project guidance and discussions. We thank Stephanie Mohr (DRSC) for guidance and assistance with whole genome screening optimization and follow-up analysis. We thank John M. Belote (Syracuse) for sharing DTS proteasome reagents and Alexey Soshnev for graphical designs. We acknowledge the *Drosophila* RNAi Screening Center (DRSC) for RNAi libraries and reagents. We acknowledge the NYU School of Medicine Genome Technology Center for high-throughput sequencing and analysis assistance. Stocks obtained from Bloomington *Drosophila* Stock Center were used in this study. We thank Toby Lieber and Lacy Barton for valuable discussion and editing of the manuscript.

---

**S1 Figure. Characterization of *Wolbachia-infected* JW18 cell line**. **(A)** Phylogenetic analysis of the *Wolbachia* strain in JW18 cells compared to previously sequenced strains (Chrostek et al., 2013). **(B)** Genome-wide copy number analysis of *Wolbachia* strain in JW18 cells. **(C)** Comparison of genome-wide copy number variation of *Wolbachia-infected* JW18 *Drosophila* cell line with *Wolbachia-free* S2R+ and Kc167 *Drosophila* cell lines. Plots of mapped DNA read density along the genome. Deduced copy number is indicated by color (see key). Genome-wide copy number analysis is shown for three *Drosophila* cell lines: *Wolbachia-infected* JW18, S2R+, and Kc167.

**S2 Figure. *Drosophila* gene expression analysis in the presence and absence of *Wolbachia***. **(A) (i)** *Wolbachia* infection in the JW18 cell line can be removed through treatment with doxycycline to generate a *Wolbachia-free* version of the cell line JW18TET. **(A)** ***(ii)** Wolbachia* infection in the JW18 cell line and the *Wolbachia-free* status of JW18TET cell line confirmed by DNA qPCR assay (see methods). **(B)** Differential gene expression analysis from RNAseq data comparing changes in host gene expression in the presence (JW18) and absence (JW18TET) of *Wolbachia.* **(C)** List of most highly upregulated (i) and most highly downregulated *(ii)* host genes in the presence of *Wolbachia* infection.

**S3 Figure. Analysis pipeline for quality control assessment and primary hit selection from the whole genome screen**. Quality control of primary data involved exclusion of outdated or non-specific dsRNA amplicons, positional effect analysis, and assessment of gene expression levels in JW18 cells. Thereafter, primary screen hits were selected based on threshold criteria and hits were categorized as low, medium, or high confidence.

**S4 Figure. Positional effect analysis on whole genome screen**. **(A)** Superimposed visual representations of the collective average robust Z scores **(*i*)** and standard deviations ***(ii)*** represented by dot sizes for each well position of all 198 384-well plates screened for the whole genome screen. RpL40 dsRNA control wells for increasing *Wolbachia* level are highlighted by the two green boxes. Doxycycline control wells for decreasing *Wolbachia* level are highlighted in two magenta boxes. Well A1 highlighted in the black box was excluded from further analysis because all 66 amplicons plated in well A1 across the screen had a very low robust Z score and the standard deviation was very high compared to all other well positions in the screen. **(B)** Visual representation of *Wolbachia* levels in all wells grouped by row **(i)** and by column **(ii)**. All visualization was done using Vortex software (Dotmatics, USA).

**S5 Figure. Whole genome screen primary results. (A)** 1117 primary hits binned into 9 bins of 124 genes each based on gene expression level from RNAseq data. * Note: Final bin only contains 123 genes. **(B)** Representation of the effect on *Wolbachia* level for primary hits within each bin (defined in A) including genes that increased (magenta) and decreased (magenta) *Wolbachia* upon RNAi knockdown. **(C)** Representation of gene DNA copy number variation of primary hits within the 9 bins (defined in **A** and **B**).

**S6 Figure. Gene Ontology analysis of whole genome screen primary results**.
Primary screen hits that increased (329 genes) *Wolbachia* levels significantly upon RNAi knockdown were analyzed for gene ontology term enrichment in biological processes, molecular processes, and cellular components. Total genes for GO term in *Drosophila melanogaster* genome shown in brackets after term. Number of genes represented shown on the bar and the number of expected genes to hit by chance shown in brackets. p-values are represented after each bar. Note: No enrichment (enrichment score >5) of any terms for screen hits that decreased *Wolbachia* levels (788 genes) was found. Gene ontology analysis was performed using PANTHERTM Version 12.0 (release 2017-07-10).

**S7 Figure. Host gene networks that influenced *Wolbachia* levels in genome-wide screen**. We identified the core ribosome (**Fig 5**), translation initiation complex (**Fig 5**), core proteasome, BRD4-pTEFb complex, Coatomer I complex, Brahma complex and components of the spliceosome as enriched for genes that increased *Wolbachia* levels in the primary screen. Three cell polarity proteins decreased *Wolbachia* levels in the primary screen. Changes in *Wolbachia* levels in the primary screen are indicated by color: increases (magenta), decreases (green), and no effect (grey). Changes in cell growth during the whole genome screen assay are indicated by icon shape: no change (circle), decrease (square), and increase (triangle). Note: These results represent the raw results from the screen prior to secondary validation.

**S8 Figure. *In vitro* and *in vivo* validation of host proteasome effect on *Wolbachia* levels**. **(A)** Validation of proteasome network by RNAi in the JW18 cell line. Representative genes were validated using dsRNA amplicons targeting unique regions of each gene. Effects on *Wolbachia* levels were assessed quantitatively by DNA qPCR measuring the number of *Wolbachia* genomes using wspB copy number relative to the *Drosophila* gene RpL11 copy number to represent host cell nuclei. Network validation is relative to untreated JW18 cells and the positive control RpL40 RNAi knockdown is included for reference. **(B)** *Wolbachia-infected* stage 10 *Drosophila* egg chamber of control sibling (TM3/TM6B) and temperature sensitive proteasome mutant sibling (DTS5/TM3) at the restrictive temperature. **(C)** Quantification of integrated density of the *Wolbachia* FISH probe in stage 10 egg chambers collected from 15-25 *Drosophila* ovary pairs for each genotype. Differences between control and mutant siblings is statistically significant (Mann Whitney, p<0.0001).

**S9 Figure. RNAseq gene expression data of core proteasome and host translation components in the presence (JW18) and absence of *Wolbachia.* (A)** The expression of genes encoding the host proteasome is upregulated in the presence of *Wolbachia* in the JW18 cell line. **(B)** The expression of host ribosome components (Rp subunits) are not different in the presence or absence of *Wolbachia.*

**S10 Figure. *Wolbachia-infected* cells do not show upregulation of stress response-related genes. (A)** RNAseq analysis on *Wolbachia-infected* JW18 cells compared to JW18TET cells highlighting stress related genes’ expression in response to *Wolbachia* infection in *Wolbachia-infected* cells. **(B, C)** JW18 cells do not show altered eIF2 phosphorylation antibody staining compared to *Wolbachia*-free cells.

## List of Supplementary Tables

S1 Table. List of outdated dsRNA amplicons that do not have gene targets.

S2 Table. List of all dsRNA amplicons that target multiple genes in whole genome screen library.

S3 Table. List of excluded hits with dsRNA amplicons that had multiple gene targets.

S4 Table. List of dsRNA amplicons seeded into well A1 in library.

S5 Table. List of genes with amplicons that hit in the screen but were excluded for no gene expression in RNAseq dataset.

S6 Table. List gene hits from whole genome RNAi screen

S7 Table. Hit dsRNA amplicons comparison of changes in Wolbachia levels and changes in cell growth.

